# Modulation of hippocampal network oscillation by PICK1-dependent cell surface expression of mGlu3 receptors

**DOI:** 10.1101/2021.11.19.469240

**Authors:** Pola Tuduri, Nathalie Bouquier, Benoit Girard, Enora Moutin, Maxime Thouaye, Julie Perroy, Federica Bertaso, Jeanne Ster

## Abstract

mGlu3 receptors control the sleep/wake architecture which plays a role in the glutamatergic pathophysiology of schizophrenia. Interestingly, mGlu3 receptors expression is decreased in the brain of schizophrenic patients. However, little is known about the molecular mechanisms regulating mGlu3 receptors at the cell membrane. Subcellular receptor localization is strongly dependent on proteinprotein interactions. Here we show that mGlu3 interacts with PICK1 and that their binding is important for receptor surface expression and function. Disruption of their interaction via an mGlu3 C-terminal mimicking peptide or an inhibitor of the PDZ domain of PICK1 altered the functional expression of mGlu3 receptors. Consequently, we investigated whether disruption of the mGlu3-PICK1 interaction affects hippocampal theta oscillations *in vitro* and *in vivo*. We found a decreased frequency of theta oscillations in organotypic hippocampal slices, similar to what previously observed in mGlu3 ^−/−^ mice. In addition, hippocampal theta power was reduced during REM sleep, NREM sleep and wake states after intra-ventricular administration of the mGlu3 C-terminal mimicking peptide. Targeting the mGlu3-PICK1 complex could thus be relevant to the pathophysiology of schizophrenia.

## Introduction

As the major neurotransmitter in the brain, glutamate is considered critical for CNS function. Indeed, a number of findings implicate glutamate as an essential contributing factor for theta oscillations [1, 2] and abnormalities in glutamatergic neurotransmission lead to altered network oscillations. Evidence indicates that dysfunctional neural oscillations represent an endophenotype of schizophrenia. Glutamate regulates the central nervous system through the actions of ionotropic (NMDA, AMPA and kainate receptors) as well as metabotropic receptors (mGlu). Efforts have been focused to characterize the clinical implications of targeting these receptors. For example, inhibiting NMDA receptor signaling induces a syndrome in healthy individuals that includes symptoms associated with schizophrenia [3]. However, excessive direct activation of NMDA receptors induces epileptic seizures and excitotoxicity-induced neuronal death [4] and cannot be considered as a safe therapeutic avenue. The focus of research has therefore shifted towards a subtler modulation of NMDA receptor function in order to reduce the risk of side effects. mGlu receptors finely modulate synaptic transmission by a variety of second messengers. Thus, they represent attractive alternative therapeutic targets for the treatment of psychiatric disorders. In particular mGlu2/3 receptors, and more specifically mGlu3, have been proposed as potential targets for schizophrenia, Parkinson’s disease, drug addiction, anxiety [5-7].

The metabotropic glutamate receptors mGlu2/3 are broadly expressed in the central nervous system where they are involved in a variety of physiological and pathophysiological processes. They control synaptic plasticity including long-term potentiation (LTP) and long-term depression (LTD) actingt mainly at presynaptic sites [8, 9]. In parallel, they also play important postsynaptic functions. mGlu2/3 share most of their pharmacology and they have been studied as a homogeneous receptor group. However, in recent years, increasing evidence suggests that postsynaptic mGlu3 receptors display a distinct localization and physiological functions in the CNS compared to the closely related mGlu2 receptor. Indeed, postsynaptic mGlu3 receptors induce a LTD of excitatory transmission onto prefrontal cortex pyramidal cells, which is prevented by stress [10]. We have previously shown that they are involved in the modulation of theta rhythms (6-14 Hz) in the CA3 area of hippocampus, which is altered in a mouse model exhibiting a schizophrenic phenotype [11]. In the hippocampus, activation of mGlu2/3 promotes the induction of synaptic plasticity by modifying postsynaptic NMDA receptor (NMDAR) function [12, 13]. mGlu2 receptors modulate primarily extrasynaptic NMDA receptors while mGlu3 receptors act on both extrasynaptic and synaptic NMDARs. These data are likely to reflect the differential somatodendritic localization of these receptors, with mGlu3 receptors being located to synaptic active sites. Despite this evidence, the molecular mechanism by which mGlu3 receptors participate in the control of these postsynaptic physiological processes remains largely undetermined. Similarly, little is known about the molecular events that take place downstream of receptor activation. Interacting proteins are associated with different domains of the receptor, in particular its C-terminal domain. These interacting proteins form a functional scaffold, or “receptosome”, that regulates the cellular targeting and specify their coupling to different signal pathways [14]. For example, the interaction between mGlu1/5 and Homer proteins plays an important role in shaping dendritic spines, suggesting a role in synaptogenesis [15] and plasticity [16] as a safeguard of brain physiology [17]. Identifying which proteins interact specifically with mGlu3 receptor and the role of these interactions are essential steps to understand physiological functions of this receptor in the neuronal network. Interestingly, yeast 2-hybrid studies have identified PICK1 [18], GRIP [18] and PP2C [19] as specific mGlu3-interacting partners. Among them, PICK1 (Protein Interacting with C Kinase 1) is a promising candidate to study the specific localization and function of mGlu3 receptor. PICK1 is expressed at high levels in brain and is localized at the neuronal synapses. The PDZ domain of PICK1 binds to a large number of proteins and its main function is to regulate the subcellular localization and surface expression of its PDZ-binding partners.

We hypothesize that the binding of PICK1 to mGlu3 receptors could be important for the localization of mGlu3 and might modulate its physiological functions, including hippocampal rhythms.

## Results

### The mGlu3 receptor interacts with PICK1

To characterize the interaction between PICK1 and mGlu3 receptor, we first used a coimmunoprecipitation approach. We transiently expressed GFP-PICK1 in the absence or presence of mGlu3 receptor bearing an HA tag in the extracellular N-terminal domain of the receptor (HA-mGlu3) in HEK-293 cells and performed co-immunoprecipitation experiments using a GFP nanobody (Figure 1A). We detected bands of stronger intensity around 200 kDa /100 kDa corresponding to mGlu3 dimers / monomers in presence of the full-length receptor and GFP-PICK1, compared to cells expressing HA-mGlu3 and GFP. As PICK1 contains a single PDZ domain that could interact with the PDZ ligand motif of mGlu3 receptors, we generated a truncated mGlu3 mutant lacking its “-SSL” PDZ ligand motif, HA-mGlu3ΔPDZlig. Deletion of the last three amino acids in the distal C-terminal of mGlu3 receptors (HA-mGlu3ΔPDZlig) almost completely prevented the interaction between mGlu3 and PICK1 (Figure 1A). This result indicates that the PDZ-ligand domain of mGlu3 receptor is necessary for its interaction with PICK1. We confirmed this interaction by performing a reverse immunoprecipitation using anti HA-antibody (Figure 1B). Given the high sequence homology of mGlu2 and mGlu3 receptor C-terminal sequences, we addressed the specificity of the interaction of PICK1 with mGlu3 receptor by performing a similar co-immunoprecipitation approach as in Fig.1A using an HA-tagged mGlu2 (HA-mGlu2) construct (Figure 1C). No band was detected in cells transfected with HA-mGlu2 and GFP-PICK1, indicating that PICK1 interacts with mGlu3 but not mGlu2.

**Fig. 1:**
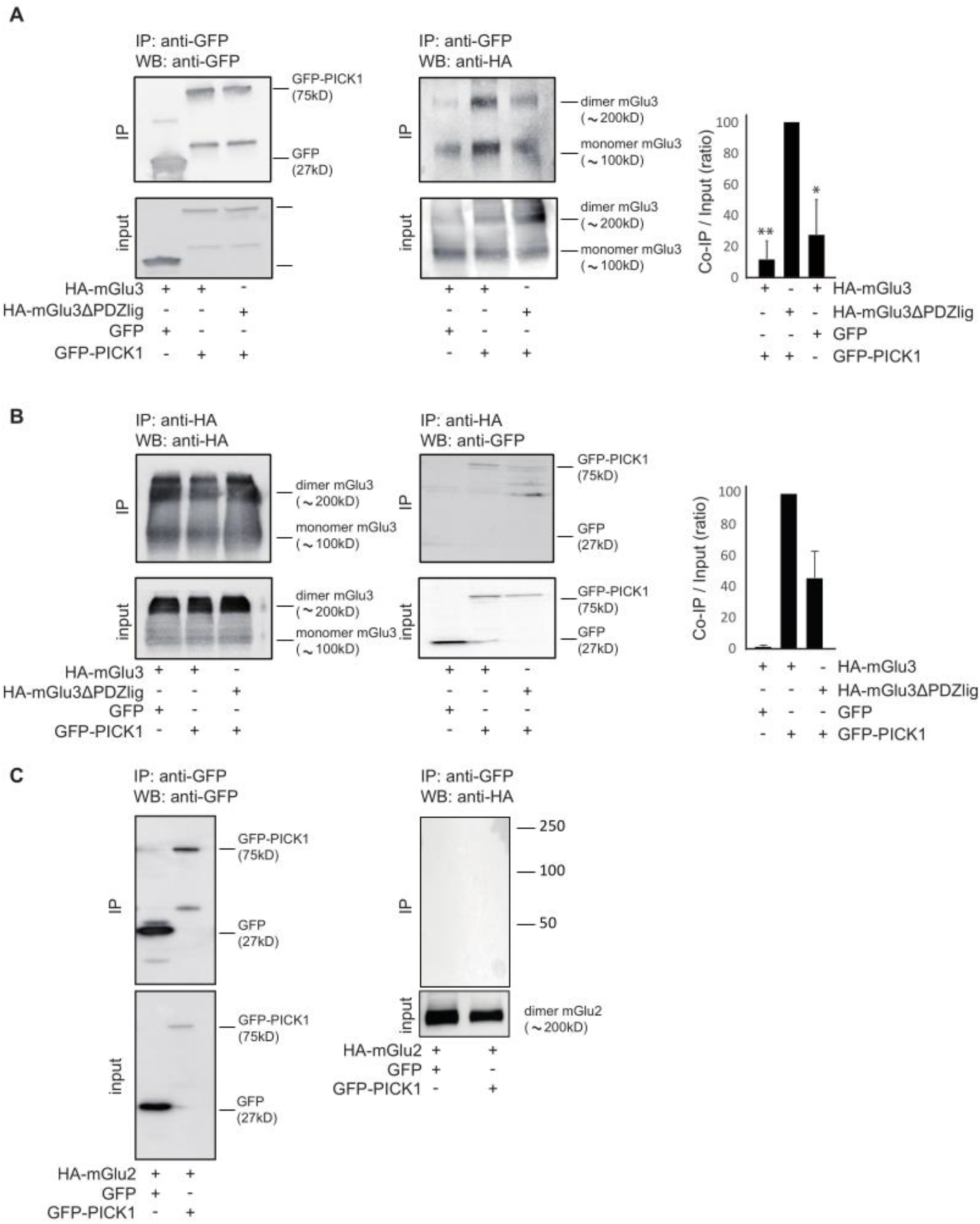
Co-immunoprecipitation of HA-mGlu3 receptor with GFP-PICK in HEK-293 cells. *Legend Figure 1/Specific interaction of HA-mGlu3 receptor with GFP-PICK1 in HEK-293 cells*. HEK-293 cells were transfected with plasmids encoding either HA-mGlu3 or HA-mGlu3ΔPDZlig and co-transfected with GFP-PICK1 or GFP. Proteins were immunoprecipitated with either GFP-Trap® beads (**A**) or monoclonal anti-HA antibody (**B**). mGlu3 receptor and PICK1 expression in inputs and immunoprecipitates were analyzed by Western blotting using anti-HA and anti-GFP antibody respectively. The histogram represents the amount of mGlu3/PICK1 binding (IP/ Input ratio). Results are means ± SEM for densitometry analyses of blots obtained in three independent experiments performed on different sets of cultured cells. **C**, HA-tagged mGlu2 receptors co-expressed with GFP-PICK1 or GFP alone were immunoprecipitated using anti-HA agarose antibody and detected using an anti-GFP antibody. Panel C is representative of two independent experiments. Mw, molecular mass (indicated in kDa). IP, immunoprecipitation.

To further demonstrate a direct interaction between mGlu3 and PICK1, we carried out pull-down experiments of cytosolic PICK1 with synthetic peptides encompassing the C-terminal domain of the mGlu3 receptor conjugated to Sepharose beads. Two different synthetic peptides were generated: one corresponding to the last 60 amino acids in the C-terminal of mGlu3 receptor (mGluR3ctWT) and a similar peptide in which the C-terminal leucine was replaced by a glutamate (mGluR3ctSSD), to hinder the interaction. Lysates of cells expressing or not PICK1 were incubated with these peptides (Figure 2A). As expected, we detected a stronger interaction with GFP-PICK1 when using mGlu3ctWT beads (~75 kDa band, detected with both anti-GFP and anti-PICK1 antibodies). We then tested the mGluR3ctWT and mGluR3ctSSD beads on mouse brain lysates (Figure 2B). The mGlu3 C-terminal co-immunoprecipitated with the native PICK1 (50 kD) expressed in mice brain, indicating that they may form a complex. We also found that the mutant bearing a point of mutation in the ligand PDZ domain (mGlu3RctSSD) partially altered the interaction with PICK1. Altogether, these results show that mGlu3 interacts specifically via its C terminal to the PDZ domain of endogenous PICK1.

**Fig. 2:**
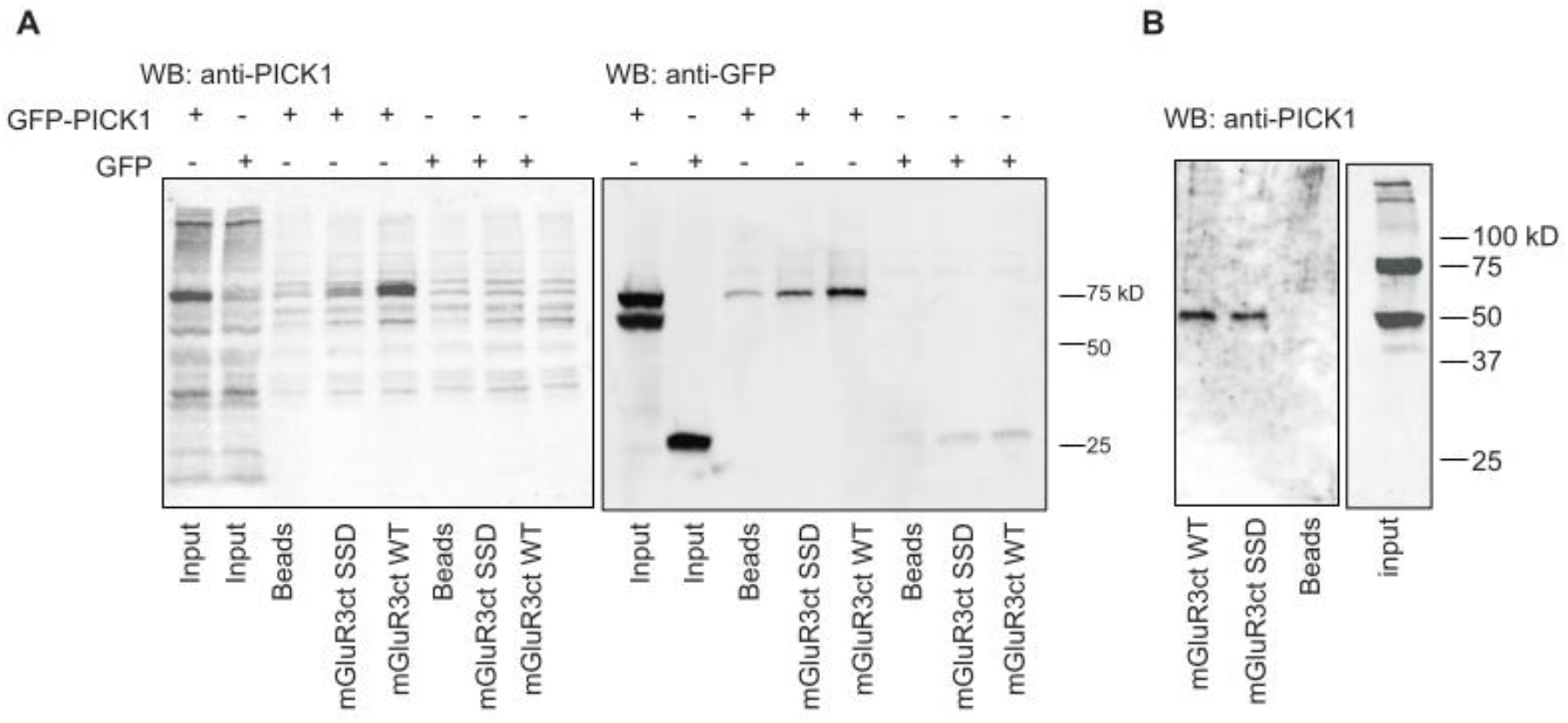
The mGlu3 C-terminal associates with native PlCK1. *Legend Figure 2/ The mGlu3 C-terminus associates with native PICK1*. **A**, HEK lysates expressing GFP-PICK1 or GFP were incubated with Sepharose-immobilized synthetic peptides that incorporated either the 20 C-terminal residues of the WT (mGlu3RctWT) or mutated mouse mGlu3 receptor (mGlu3ctSSD) or with Sepharose beads only. **B**, Protein extracts from mouse brain were incubated with the indicated Sepharose-immobilized peptides, and bound proteins were analyzed by Western blotting with an anti-PICK1 antibody. Note that native PICK1 expressed in mice brain co-immunoprecipitates with the mGlu3 receptor C-terminal. Representative data of three independent experiments are illustrated.

### Peptidic and pharmacological tools to uncouple binding of mGlu3 to PICK1

To determine the functional role of the mGlu3-PICK1 coupling, we used and engineered tools to disrupt their interaction. First, we designed a competitive peptide encompassing the last ten C-terminal amino acids of the mGlu3 receptor, which included the PDZ ligand motif SSL, conjugated to the cell-membrane transduction domain of the HIV-1 Tat protein (TAT-mGlu3, Figure 3A). A control peptide was prepared bearing the same overall sequence with the exception of the last 3 amino acid of the PDZ ligand motif, SSL, which were mutated to alanine (AAA). To evaluate the binding of PICK1 to mGlu3 receptors in presence of the TAT-mGlu3, we used a co-immunoprecipitation approach as Fig.1A (Figure 3B). There was a strong reduction in the intensity of band corresponding to the receptor (around 200 kDa) in cells co-transfected with HA-mGlu3 and GFP-PICK1 in presence of TAT-mGlu3 compared to TAT-control.

**Fig. 3:**
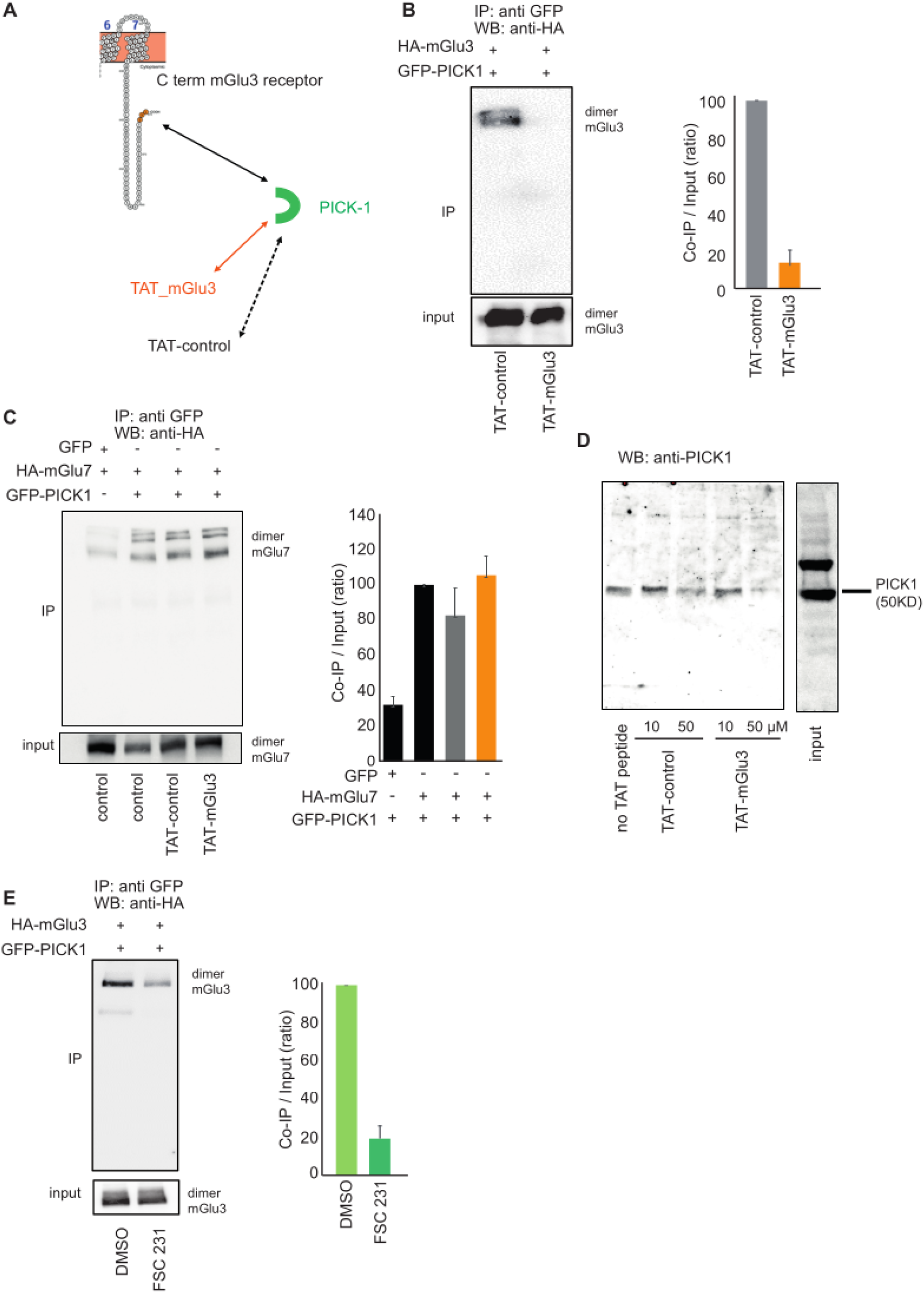
Pharmacological tools uncoupling mGlu3-PICK1 interaction *in vitro*. *Legend Figure 3/* Pharmacological tools uncoupling mGlu3 - PICK1 interaction *in vitro*. **A**, schematic representation of the action of TAT peptides on PDZ interaction with mGlu3 C-terminal. Co-immunoprecipitation of HA-mGlu3 (anti-HA antibody) and PICK1 from HEK-293 lysates with either no drug, with TAT peptides (10 μM each; **B**), FSC231 (25 μM) or DMSO (25 μM; **E**). HEK-293 cells were co-transfected with plasmids encoding HA-mGlu3 and GFP-PICK1. Proteins were immunoprecipitated with GFP-Trap® beads. Total protein extracts were analyzed by Western blotting using anti-HA antibody. **C**, HA-tagged mGlu7 receptor co-expressed with GFP-PICK1 or GFP alone was immunoprecipitated using agarose conjugated anti-HA antibody and detected using an anti-HA antibody. **D**, Pull-down experiment carried out using mouse brain extracts supplemented with no TAT peptide, 10 or 50 μM TAT-control peptide and 10 or 50 μM TAT-mGlu3 peptide. **B**, **C**, **D**, **E** are representative of three independent experiments. **B**, **C**, **E**, the histograms represent the amount of mGlu3-PICK1 binding. Results are means ± SEM for densitometry analyses of blots obtained in three independent experiments performed on different sets of cultured cells.

To confirm the specificity of the TAT-mGlu3 to mGlu3 receptor, we performed a similar co-immunoprecipitation targeting the mGlu7 receptor, which also interacts with PICK1 [18], Figure 3C). We observed that binding of PICK1 to the mGlu7 receptor was prevented by neither peptide. The specificity of the TAT peptides was further characterized by pull-down experiments. We detected a dose-dependent decrease in native PICK1 binding to mGlu3 only when TAT-mGlu3 was added to the mouse brain lysate (Figure 3D). Taken together these data confirmed the specific competition between TAT-mGlu3 and mGlu3 for its receptosome. The second strategy was to perform co-immunoprecipitation experiments using a small PICK1 PDZ-binding molecule, FSC231, which is known to prevent PICK interactions [20]. This compound altered the binding of PICK1 to mGlu3 receptor compared to the control (DMSO, Figure 3E).

### PICK1 regulates mGlu3 receptor surface expression via the PDZ ligand motif

Membrane proteins localization depends on interactions with intracellular binding partners. We tested whether this was the case for mGlu3 receptor and PICK1, at first by using a heterologous expression system, HEK-293 cells. We monitored the constitutive trafficking of an HA-tagged version of the receptor together with either GFP-tagged PICK1 (GFP-PICK1) or GFP (Figure 4A). We visualized HA antibody-labeled surface receptors before permeabilization (Alexa Fluor 350 tagged secondary antibody) and internalized receptors after permeabilization (Cy3 tagged secondary antibody). When transfected with GFP, the receptor was diffusely distributed throughout the cell (ratio surface *vs.* cytoplasm = 1.64 ± 0.08). Internalized mGlu3 receptors formed clusters in the cytoplasm. By contrast, when the receptor was transfected with GFP-PICK1, surface expression of HA-mGlu3 was significantly increased (ratio surface *vs.* cytoplasm = 2.32 ± 0.28). Moreover, most of the internalized receptors were located near the cell membrane. These results indicated that the localization of mGlu3 receptors depends on PICK1 expression in HEK-293 cells.

**Fig. 4:**
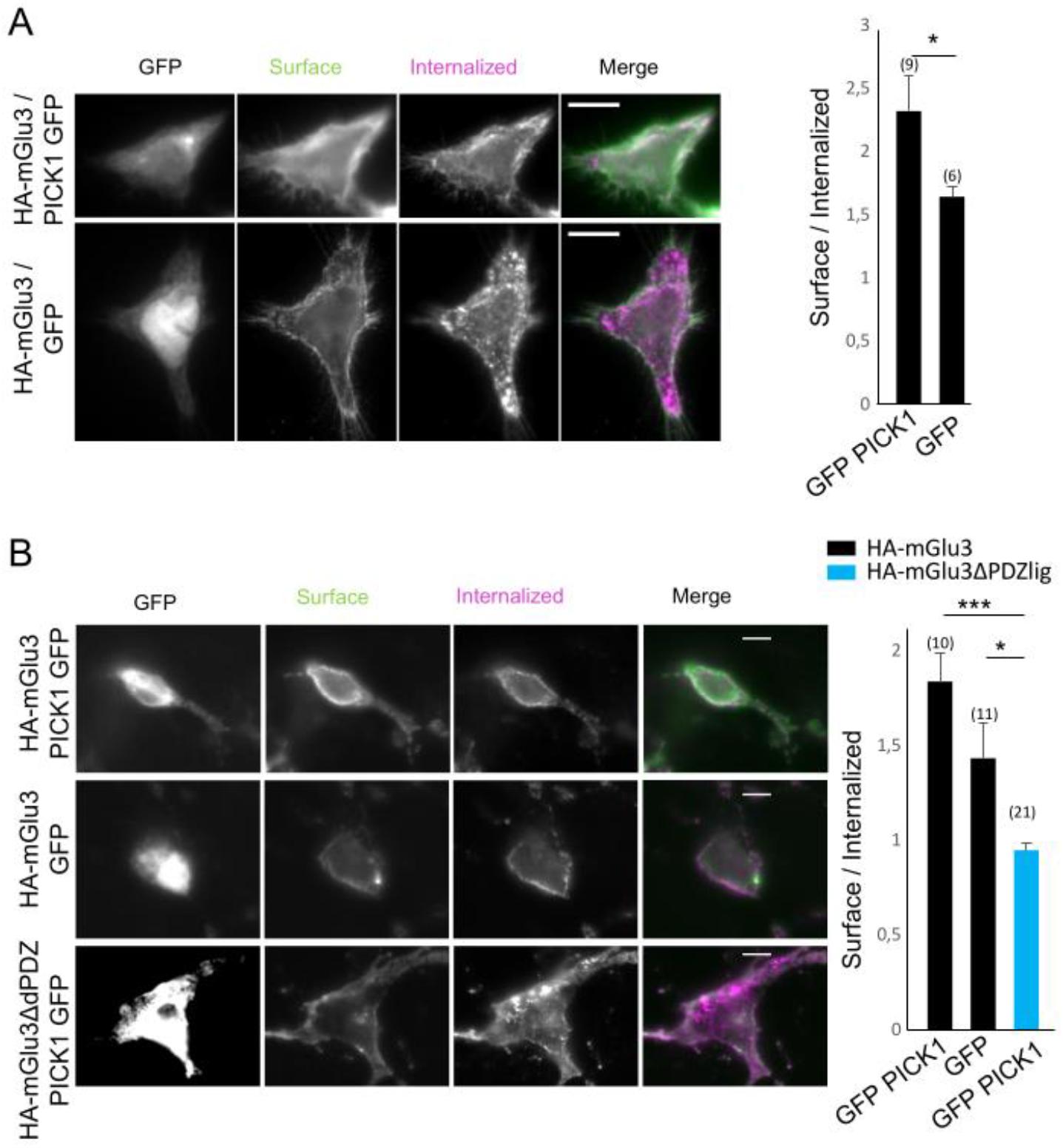
mGlu3 receptor internalization is regulated by PICK1. *Legend Figure 4/ mGlu3 receptor internalization is regulated byPICK1*. **A**, HEK-293 cells expressing wild-type mGlu3 receptor in presence or absence of PICK1. mGlu3 receptors were labeled with an anti-HA antibody (pink, internalized proteins, green, surface-expressed receptor). Summary histogram quantifying the internalization of mGlu3 receptors. **B**, Hippocampal neurons were transiently co-transfected with HA-mGlu3 and GFP-PICK1 or with HA-mGlu3ΔPDZlig and GFP-PICK1. Summary histogram quantifying internalization of mGlu3 receptors. Data represent means ± SEM; *p< 0.05, ***p < 0.001. Scale bars =10 μm.

To deepen this characterization, we studied the localization of mGlu3 and PICK1 in primary hippocampal neurons and found similar results to those observed in HEK-293 cells (Figure 4B, 4C). In neurons co-transfected with HA-mGlu3 and GFP-PICK1, surface levels of mGlu3 receptors were higher than in neurons transfected with HA-mGlu3 alone. Previous studies have shown that the PDZ ligand residues of the intracellular C terminal of mGlu receptors are critical for their synaptic localization [21]. We tested whether the PDZ ligand also plays a role in mGlu3 receptor endocytosis. The surface expression of the HA-mGlu3ΔPDZlig receptor lacking the PDZ ligand motif was significantly decreased compared to the wild type receptor (Figure 4C; ratio surface / internalized: GFP-PICK1 / HA-mGlu3 = 1.84 ± 0.15; GFP / HA-mGlu3 = 1.43 ± 0.18; GFP-PICK1 / HA-mGlu3ΔPDZlig = 0.95 ± 0.04). Altogether, these results show that PICK1 promotes the expression of mGlu3 receptors at the neuronal membrane.

### Binding to PICK1 is essential for the functional expression of the mGlu3 receptor

We reasoned that an alteration of mGlu3-PICK1 interaction could alter the subcellular localization of the receptors. We examined the effect of an acute treatment with TAT-mGlu3 (1 hour at 1 μM; Figure 5A, B) or FSC231 (1 hour at 25 μM; Figure 5C, D) on the subcellular localization of mGlu3 receptors in neurons. Both treatments induced a significant reduction in mGlu3 receptor surface expression (Figure 5; ratio surface / internalized: TAT-control = 1.16 ± 0.06; TAT-mGlu3 = 1 ± 0.03; DMSO = 1 ± 0.03; FSC231 = 0.84 ± 0.06). Conversely, no significant effect was induced by the TAT-control peptide (Figure 5). Next, to determine the physiological significance of the mGlu3-PICK1 interaction, we performed patch-clamp recording of CA3 pyramidal cells (PCs) and interneurons in organotypic hippocampal slice cultures in the presence of TAT-mGlu3 peptide or FSC231. As previously reported [22], the activation of mGlu3 receptors by LCCG-1 (10 μM) induced a significant inward current in CA3 pyramidal cells (PCs) which is obvious under conditions of synaptic transmission block (TTX /DAP5; Figure 6A). The LCCG-1-induced inward current was blocked in presence of TAT-mGlu3 (10 μM) (−6.63 ± 2.89 pA, *p<0.01, Figure 6A, B) in CA3 PCs as well as in interneurons (−2.1 ± 1.86 pA, **p<0.01, Figure 6C). The TAT-control peptide had no effect in PCs (10 μM, CA3 PCs: control=-23.29 ± 1.00 pA, TAT-control = −23.86 ± 3.56 pA) and interneurons (CA3 interneuron: control = −21.75 ± 4.60 pA, TAT-control = −15 ± 1.93 pA; Figure 6A, B, C). Similar results were obtained in presence of the PDZ domain inhibitor FSC231: the LCCG-1-induced inward current was blocked in CA3 PCs and interneurons (PCs: DMSO = −27.00 ± 4.40 pA, FSC231 = 5.86 ± 3.48 pA, **p<0.01; CA3 interneurons: DMSO= −20.76 ± 5.66 pA, FSC231 = 2.10 ± 2.00 pA, **p<0.01; Figure 6A, B, C). Moreover, the spontaneous synaptic activity is not altered by treatment with the TAT peptides, suggesting that AMPA function is preserved (Figure 6D; control = −28.34 ± 4.83 pA, TAT-control = −24.12 ± 5.33 pA; TAT-mGlu3 = −25.26 ± 3.98 pA).

**Fig. 5:**
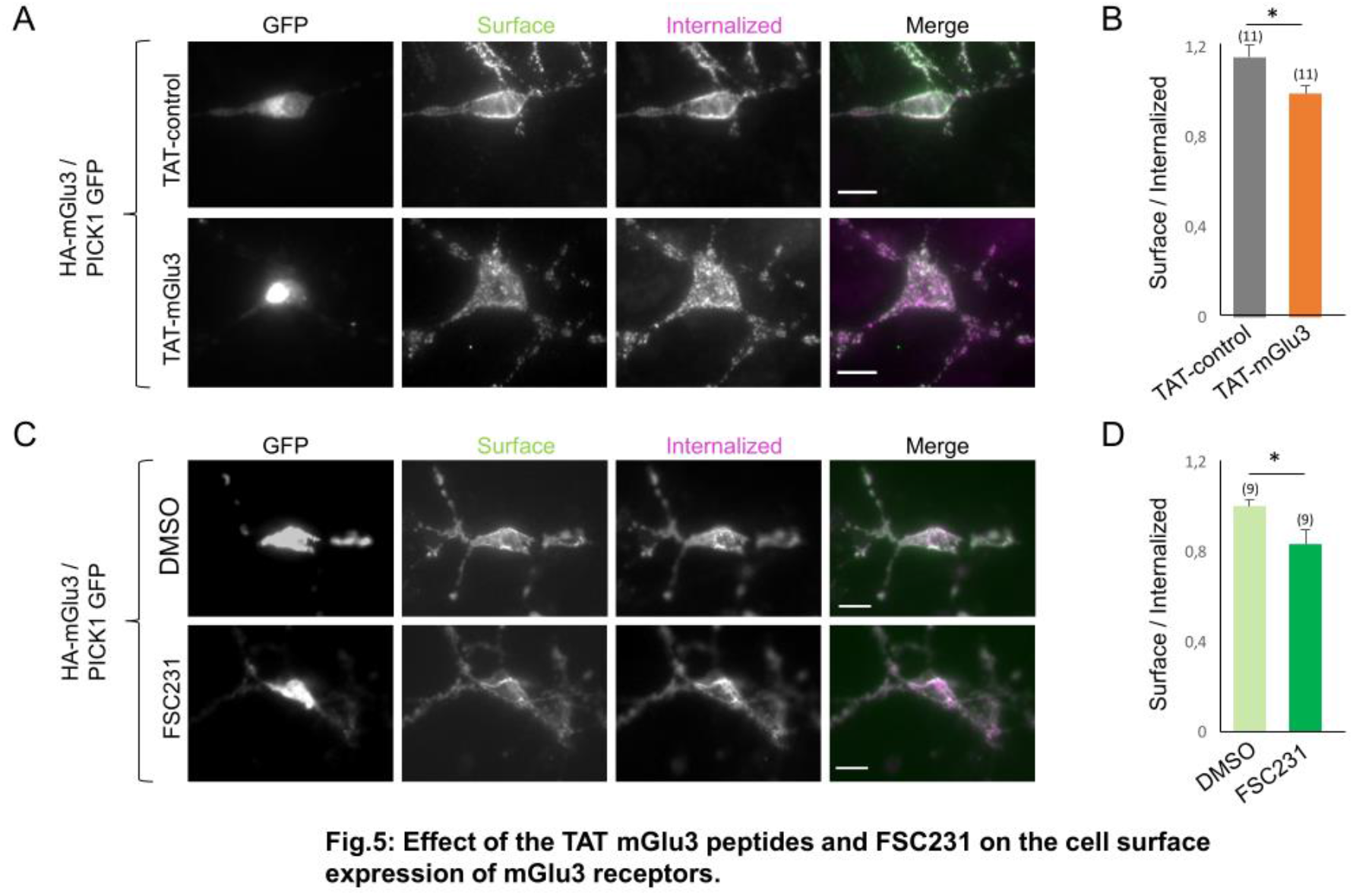
Effect of the TAT mGlus peptides and FSC231 on the cell surface expression of mGlus3 receptors. *Legend Figure 5/ Effect of the TAT-mGlu3 peptide or FSC231 on the cell surface expression of mGlu3 receptors*. **A**, **B** Hippocampal neurons were transiently co-transfected with HA-mGlu3 and PICK1. Neurons were treated with DMSO alone, FSC231, TAT-mGlu3 or TAT-control peptides at 37°C for 1 hour. Then neurons were labelled with anti-HA antibody, washed and returned to conditioned media containing the same treatment at 37° for 15 min. The cells were stained and images acquired as described in Fig.4. Scale bar, 10 μm. **C**, summary histogram quantifying the internalization ratio from panel A and B. Data represent means ± SEM; *p< 0.01, ***p< 0.001.

**Fig. 6:**
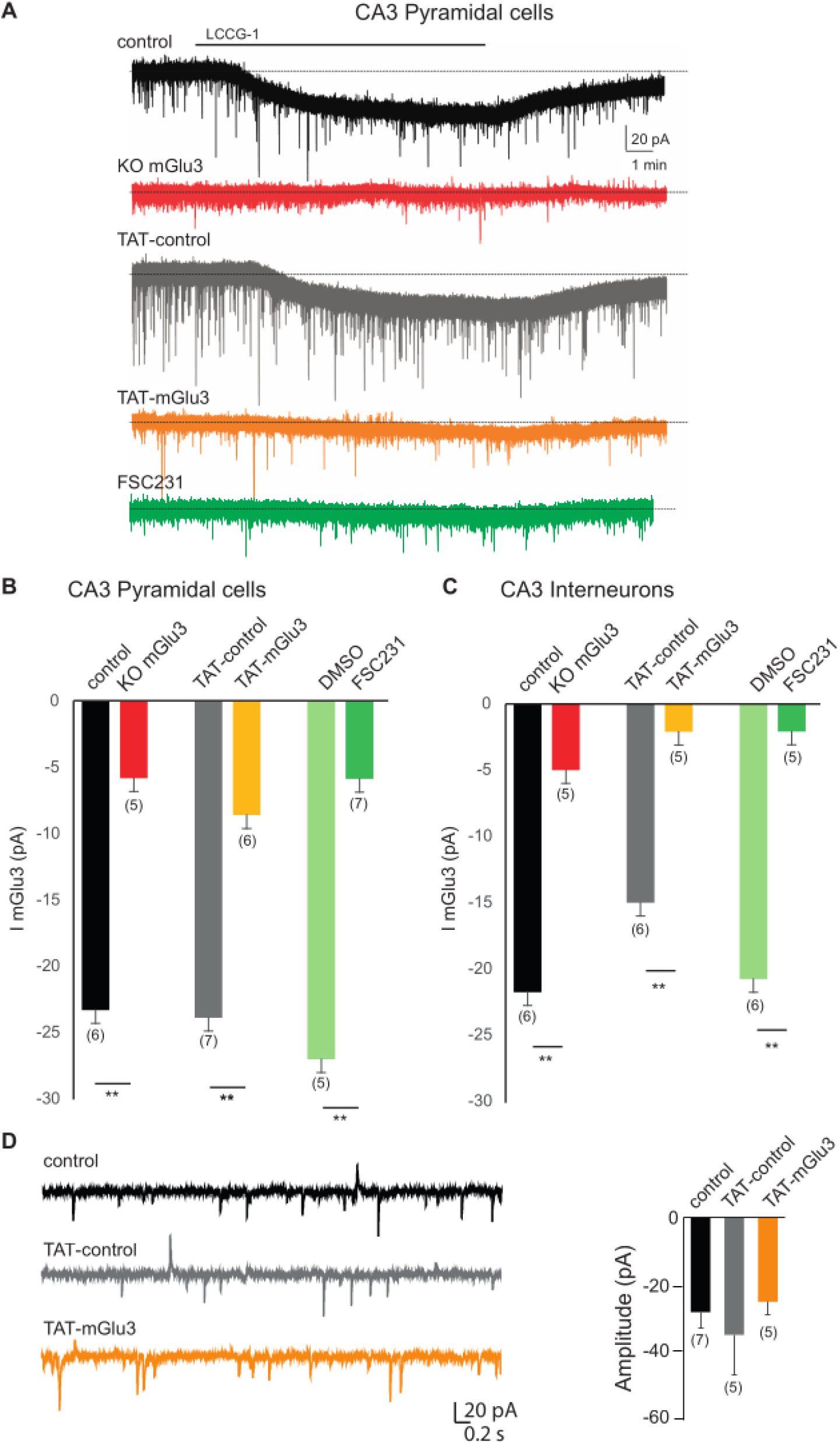
TAT-mGlus3 peptide or FSC231 specifically aitered mGlus3 receptor function in hippocampal organotypic slices. *Legend Figure 6/ TAT-mGlu3 peptide or FSC231 disturbed the functional mGlu3 receptor specifically in hippocampal organotypic slices*. **A**, Bath application of the LCCG1 in presence of TTX (1 μM), picrotoxin (100 μM), and D-AP5 (40 μM) induces inward current in voltage-clamped CA3 PCs with or without TAT-control (grey trace, black trace respectively). The inward current is not detected in CA3 PCs in presence of TAT-mGlu3 peptide (orange trace) or FSC231 (dark green trace). Histograms represent the mean ImGlu3 current to illustrate the effect of TAT-mGlu3 peptides or FSC231 in PCs (**B**) and in interneurons (**C**). **D**, Representative activity recorded in a CA3 PCs (voltage-clamped at −70mV) in presence of TAT-mGlu3 peptides or FSC231. Data represent means ± SEM; *p< 0.01, ***p< 0.001

### Theta oscillations in the CA3 area of the hippocampus are altered by impairing the mGlu3-PICK1 interaction

Cholinergic activation of CA3 network in the hippocampus results in cycles of rhythmicity [23, 24]. The mGlu3-dependent inward current contributes to the theta oscillations in hippocampal organotypic slices [22]. As we have shown that this current is also dependent on PICK1 biding to the receptor (Figure 6), we hypothesized that the mGlu3-PICK1 complex could modulate the hippocampal network activity. Theta oscillations were induced in organotypic hippocampal slices by applying the muscarinic receptor agonist methacholine (50 nM MCh, 20 min; frequency = 12.9 ± 0.38 Hz, episodes / min = 10.8 ± 2.60) [25]. The frequency of theta oscillations was significantly reduced in presence of the TAT-mGlu3 peptide compared to TAT-control (frequency: TAT-mGlu3 = 9.97 ± 0.42 Hz, TAT-control = 12.64 ± 0.75 Hz, n = 6, **p<0.01; Figure 7A, B). A similar reduction in theta frequency was observed when applying FSC231 (frequency: DMSO = 14.15 ± 0.64, DMSO / FSC231 = 10.54 ± 1.11, *P < 0.05; Figure 7A, B). The occurrence of theta episodes was not significantly altered in presence of these treatments (episodes / min: TAT-mGlu3 = 11.40 ± 2.60, TAT-control = 10.33 ± 1.67; DMSO = 11 ± 4.44, DMSO / FSC231= 7.8 ± 1.98; Figure 7C). Altogether, these results suggest: 1) a contribution of mGlu3-PICK1 to hippocampal theta oscillations in CA3 area and 2) no role in the generation of hippocampal theta rhythms.

**Fig. 7:**
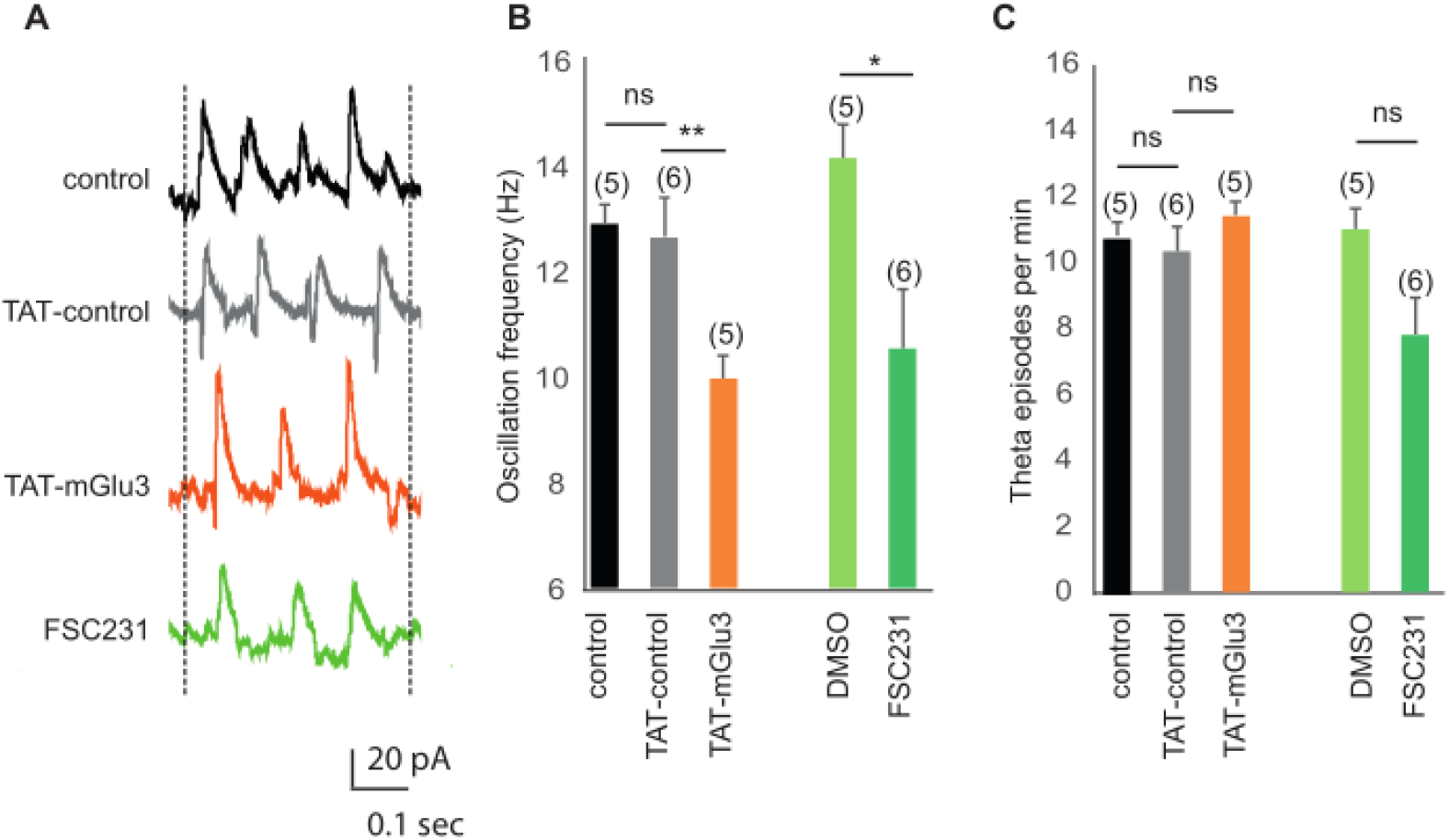
Methacholine-induced oscillations in the CA3 network are altered when PICK1 binding ro mGlu3 is disrupted. *Legend Figure 7/ Metacholine-induced oscillations in the CA3 network are altered when PICK1 binding to mGlu3 is disrupted*. **A**, Representative traces of methacholine-induced theta oscillation recorded in CA3 pyramidal cells voltage clamped at −70mV in control condition (black trace) and in presence of TAT-control (grey trace), TAT-mGlu3 (orange trace) or FSC231 (green trace). **B**, theta frequency was significantly decreased by the incubation with TAT-mGlu3 (10 μM, 1 hour incubation), but not by the incubation of TAT-control. The FSC231 (25 μM, 30 min) also reduced the theta frequency similarly to the TAT-mGlu3. **C**, The occurrence of theta episodes was much lower in presence TAT-mGlu3 or FSC231 than in presence TAT-control or DMSO.

### Effect of TAT-mGlu3 peptide on theta rhythms during sleep / wake states

During rapid eye movement sleep (REMs) and wake periods of increased attention in mice and rat, theta oscillations are observed in local field potential recordings from cortical structures, including the hippocampus [23, 26]. We asked what is the impact of the mGlu3-PICK1 complex on theta rhythms during sleep / wake states *in vivo*. We performed EEG recordings in mice injected with the TAT peptides via a cannula placed in the lateral ventricle. We confirmed the diffusion of the peptides in the hippocampus by detecting a fluorescent TAMRA-tagged version of TAT-mGlu3 35 minutes postinjection (Figure 8A). Note that no peptide was detected in other regions, including the cortex. We tested the effect of the TAT-mGlu3 during sleep / wake states in freely behaving mice. By using a sleep scoring procedure based on movement detection and electromyogram (EMG) signal analysis, we distinguished three different states: wake, non-REM sleep (NREM) and REM sleep (Figure 8B). We focused on the period between 10 and 35 minute post-injection as the optimal time for peptide diffusion. The injection of TAT-mGlu3 resulted in significant reduction of REM sleep theta power measured in the dorsal hippocampal CA3 area (Figure 8B; relative power low theta: baseline = 3.33 ± 0.33, post TAT-control = 3.45 ± 0.15, post TAT-mGlu3 = 2.25 ± 0.18; relative power high theta: baseline = 1.24 ± 0.09, post TAT-control = 1.52 ± 0.19, post TAT-mGlu3 = 0.97 ± 0.07). A decrease of theta power was also observed during NREM sleep (Figure 8B; relative power low theta: baseline = 2.59 ± 0.12, post TAT-control = 2.35 ± 0.36, post TAT-mGlu3 = 1.96 ± 0.15; relative power high theta: baseline = 1.35 ± 0.15, post TAT-control = 1.4 ± 0.2, post TAT-mGlu3 = 1 ± 0.16) and wake states (Figure 8B; relative power low theta: baseline = 2.85 ± 0.24, post TAT-control = 3 ± 0.34, post TAT-mGlu3 = 1.91 ± 0.21; relative power high theta: baseline = 1.22 ± 0.08, post TAT-control = 1.17 ± 0.1, post TAT-mGlu3 = 1.11 ± 0.17). No other frequency band was affected and the spectra profile returned to the baseline less than 1 hour after peptide injection. The TAT-control peptide had no detectable effect on the EEG recordings.

**Fig. 8:**
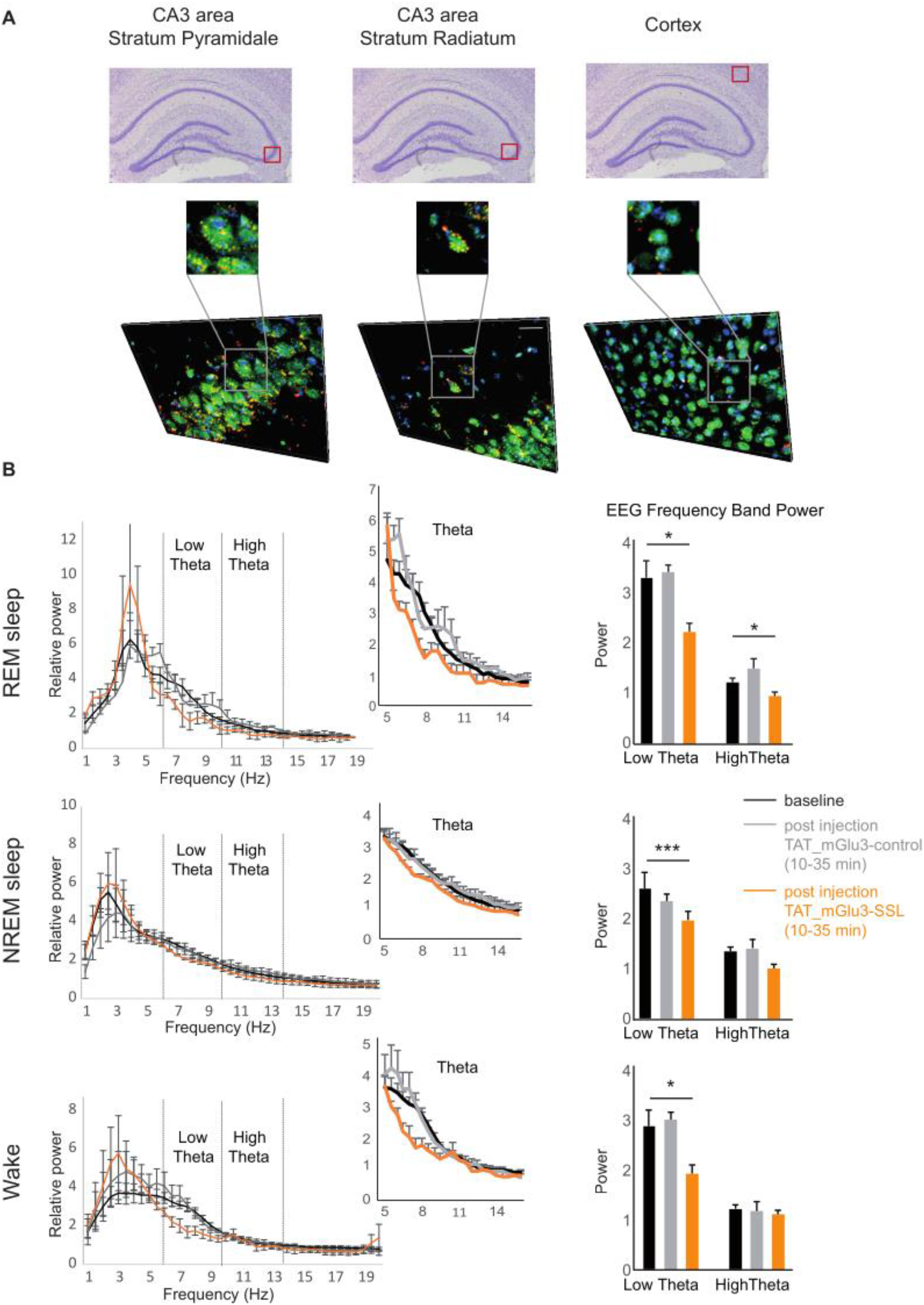
Effect of TAT mGlu3-SSL peptide on theta rhythms during sleep/wake states. *Legend Figure 8/ Effect of TAT-mGlu3 peptide on theta rhythms during sleep/Wake states*. **A**, TAT-mGlu3 TAMRA peptide was injected into the ventricle of C57Bl6/J adult mice. TAMRA expression was detected in hippocampal cells but not in the cortex. **B**. Effect of intra-ventricular injection of the TAT-control or TAT-mGlu3 peptides (5 μl, 500 μM, 750 nl / ml) on the modification of the power spectrum during REM sleep, NREM sleep and wake (n=5).

## Discussion

In this study, we demonstrated that the PDZ ligand domain–containing mGlu3 receptor interacts with PICK1 and that this domain is required for their physical and functional interactions both *in vitro* and *in vivo*. In addition, we showed that such interaction contributes to the targeting of receptors in neurons, and significantly modulates the hippocampal network activity *in vitro* and *in vivo*.

### mGlu3 interacts specifically with PICK1

Protein interactions mediated by PDZ-domains show great versatility, as PDZ domains bind to small C-terminal peptides (through class I, II and III binding motifs), internal protein segments, other PDZ domains or even lipids [27]. mGlu2 and mGlu3 receptors possess the same and common (S/T)x(V/L) motif “SSL” that belongs to the class I PDZ binding motifs. Yet, only the mGlu3 receptor binds to PICK1. Indeed, by using co-immunoprecipitation, we found that PICK1 binds specifically to mGlu3 receptor and not the closely related mGlu2 receptor. We found that the TAT-mGlu3 disturb the interaction between mGlu3 and PICK1. Thus, TAT-mGlu3 allows to distinguish the functions performed specifically by mGlu3 versus mGlu2. Therefore, a sequence of mGlu3 receptor outside its PDZ binding motif could contribute to the specificity of interaction with PICK1. Intriguingly, the total deletion of the last 3 aa corresponding to the PDZ-ligand domain of mGlu3 (mGlu3ΔPDZlig) impairs the interaction with PICK1 whereas a point of mutation of the last amino acid of the PDZ motif of mGlu3 receptors only reduces this interaction. These results suggest that the entire PDZ binding motif of mGlu3 receptor is required.

### The functional significance of the interaction of mGlu3 receptor and PICK1

Protein–receptor interactions influence signal transduction mechanisms, trafficking and localization of the receptor proteins [28] [29]. A large number of studies have established the importance of PDZ domain-mediated interactions in the localization and compartmentalization of several receptors and channels [30]. To date, little is known about the molecular factors that govern the regulatory mechanisms associated with cell surface expression of mGlu3 receptors. Proteins that specifically interact with the PDZ ligand domain of mGlu3 receptor have not been clearly characterized. To define whether the mGlu3-PICK1 interaction regulates receptor surface expression, we examined the localization of mGlu3 receptor in cultured neurons transfected with either a full-length or a truncated form of mGlu3, with or without PICK1. We found that removal of the PDZ ligand on mGlu3 receptor or the absence of PICK1 considerably reduces surface expression of mGlu3 receptor. These findings confirmed a key role for PICK1 in synaptic regulation, by adding the mGlu3 receptor to the list of its interacting partners. Indeed in the brain, PICK1 is expressed in both the pre- and postsynaptic elements, where it serves different roles. For example, PICK1 is a key regulator of postsynaptic GluA2 AMPA receptor during plasticity [31, 32]. At the presynapse, PICK1 regulates the synaptic localization and function of the mGlu7 receptor and of dopamine and norepinephrine transporters [33, 34].

Our electrophysiological data support the notion that postsynaptic mGlu3 receptors and PICK1 form a functional protein complex at the neuronal surface. Using the TAT-mGlu3 peptide, designed to disturb the endogenous interaction between mGlu3 and PICK1, we observed a decrease of the postsynaptic mGlu3-mediated inward current in hippocampal PCs and interneurons. Unfortunately, there is no specific antibody for mGlu3 to determine an alteration in their localization. Our data suggest that alteration of the mGlu3-PICK1 interaction reduces functional mGlu3 receptors at the neuronal cell surface and point to the physiological relevance for the interaction of postsynaptic mGlu3 receptor and PICK1. Our data do not exclude an interaction of the two proteins at the presynaptic site, where it could potentially modulate the excitatory / inhibitory neurotransmitter release.

Likewise, we cannot rule out the possibility that PICK1 could also influence and / or participate to mGlu3 signal transduction mechanisms, as it does for other glutamate receptors including GluA2 and mGlu7 [31, 33]. PICK1 bind protein kinase C α-subunit (PKCα) and mGlu3 receptors could signal through activation of a PLC-PKC-dependent pathway [12]. Additional investigation of such molecular mechanisms would be interesting.

### The physiological relevance of the mGlu3-PICK1 complex

Hippocampal theta oscillations (HTO) are prominent local field potentials occurring in the 4–14 Hz frequency range and generated essentially by the hippocampus [23]. Increasing evidence implicates group II metabotropic glutamate receptors in HTO both *in vitro* and *in vivo* [22, 35, 36]. Modulation of mGlu2 and mGlu3 receptors influences theta oscillations *in vivo* using different rat strains [37]. We have examined whether disrupting the mGlu3-PICK1 complex could affect theta rhythms. In organotypic hippocampal slices, Mch-mediated theta oscillations are decreased by treatment with the TAT-mGlu3 peptide and the PICK1 binding molecule, FSC231. *In vivo*, theta oscillations are most prominent during wake periods of increased attention and rapid-eye-movement (REM) sleep [23]. mGlu2/3-modulating drugs (agonist, antagonist) profoundly influence theta oscillation suggesting involvement of both receptor types in the control of theta oscillations [35-39]. In freely moving mice, we observed a significant reduction of the theta frequency power during REM sleep, NREM sleep and wake states upon intracerebral injection of the TAT-mGlu3 peptide. HTO control the timing of activity across neuronal populations in the hippocampus, prefrontal cortex, and amygdala and coordinate gamma oscillatory activity. Consequently, theta oscillations are suggested to be important in cognitive functions [40, 41]. Thus, even small changes in baseline HTO frequencies during REM sleep and quiet waking are likely to alter neural activity across large distributed brain networks, ultimately generating modifications in behavioral processes. Our data suggest that alteration of mGlu3-PICK1 interaction modifies HTO during sleep / wake states and could predict a deficit in learning rates.

### Opening on schizophrenia

Disruption of neural oscillations and synchrony may play an important role in the pathophysiology of schizophrenia, a neurodevelopmental disorder marked by abnormalities in sensory processing and cognition. Although recent research is primarily focusing on high frequency oscillations, there is also evidence of disturbances in slow rhythms in the delta and theta bands in schizophrenia [42-45]. To date, group II mGlu receptors draw a great interest as targets to treat such psychiatric conditions. Identification of new proteins that associate specifically to mGlu3 receptors will advance our understanding of the regulatory mechanisms associated with their targeting and function and ultimately might provide new therapeutic strategies to counter these psychiatric conditions.

## Material and Methods

### Plasmids and peptides

The pRK5 HA-mGlu3 cDNA was a kind gift of Jean-Philippe PIN (IGF, Montpellier, France). The plasmid encoding mGlu3ΔPDZlig was obtained from the pRK5 HA-mGlu3WT cDNA. The C-terminal sequence TSSL was replaced by a stop codon and a BamHI restriction site by site-directed mutagenesis using the following primers: 5′-CTGGACTCCACCTGAGGATCCTTGTGATACGCAGTTCA-3′ and TGCGTATCACAA GGATCC TCA GGTGGAGTCCAGGACTTC. The GFP-PICK1 cDNA was a kind gift of Oussama El Far (UNIS, Marseille, France). The mGlu3-SSL and –SSD peptides for pull-down experiments were synthesized by Eurogentec, with a purity > 70%. The sequence of the mGlu3-SSD peptide was AA AQN LYF QGP QKN VVT HRL HLN RFS VSG TAT TYS QSS AST YVP TVC NGR EVL DST TSS D, and the sequence of the mGlu3-SSL peptide was AA AQN LYF QGP QKN VVT HRL HLN RFS VSG TAT TYS QSS AST YVP TVC NGR EVL DST TSS L.

TAT mGlu3 peptides for *ex-vivo* and *in vivo* were synthesized with a purity > 95% (Eurogentec, Belgium) with the following sequences: TAT-control peptide: YGR KKR RQR RRE VLD AAA; TAT-mGlu3 peptide: YGR KKR RQR RRE VLD SSL. Both peptides were diluted to 1 or 10 μM in the appropriate media depending on the type of experiment.

The TAMRA-tagged TAT-mGlu3 peptide was synthesized by Smart Bioscience (France), with a purity > 95%. The sequence of TAMRA TAT-mGlu3 peptide was TAMRA-YGR KKR RQR RRE VLD SSL.

### Drugs

LCCG-1 D-AP5, methacholine chloride and TTX were purchased from Tocris Bioscience (Bristol, United Kingdom) and FSC231 from Sigma-Aldrich (France). FSC231 was first dissolved in DMSO and then added to the saline solutions.

### HEK cells/ transfection

HEK-293 cells were cultured in complemented Dulbecco’s modified Eagle’s medium (DMEM) as previously described [46]. Cells were transfected at 40-50% confluence using PEI (Polysciences Inc.) [47] and used 24 hours after transfection.

### Hippocampal primary cultures/ transfection

Cultures were prepared from WT C57Bl6/J as previously described [48]. Briefly, hippocampi were mechanically and enzymatically dissociated with papain (Sigma-aldrich) and hippocampal cells were seeded in Neurobasal-A medium (Gibco) supplemented with B-27 (Gibco), Glutamax (Gibco), L-glutamine (Gibco), antibiotics (Gibco) and Fetal Bovine Serum (Gibco). After 2 days in culture, Cytosine β-D-arabinofuranoside hydrochloride (Sigma-aldrich) was added to curb glia proliferation. The day after, 75% of the medium was replaced by BrainPhys medium (Stemcell technologies) supplemented with B-27 (Gibco), Glutamax (Gibco) and antibiotics (Gibco). Neurons are co-transfected with several plasmids (HA-mGlu3, HA-mGlu3ΔPDZlig, GFP-PICK1, GFP) by using Lipofectamine at DIV10.

### Co-immunoprecipitation experiments

HEK-293 cells were solubilized in a buffer containing 50 mM Tris/HCl, pH 7.4, 100mM NaCl, 2 mM EDTA, Triton 1%, 2mM DTT, 2mM MgCl2 and a protease inhibitor cocktail (Roche). Samples (1 mg) were incubated with GFP-Trap beads (Chromotek) or anti-HA antibody Sepharose beads (Sigma-Aldrich, A2095) for 1 hour at 4°C. Beads were washed five times with 50 mM Tris / HCl, pH 7.4, 250 mM NaCl, 2 mM EDTA, Triton 1%, 2mM DTT, 2mM MgCl2 and immunoprecipitated proteins were analyzed by Western blotting.

### Pull-down experiment

The mGlu3-SSL or –SSD peptides were coupled to activated CnBr-sepharose 4B (GE-Healthcare) according to the manufacturer’s instructions. Hippocampi from C57BL/6 mice (Janvier Labs) were extracted, homogenized in lysis buffer containing (in mM) 50 Tris-HCL (pH 7.4), 0.5 EDTA (pH 7.4), 1.3% CHAPS (Euromedex), protease and phosphatase inhibitor cocktails (Roche) using a potter and then centrifuged at 10,000 g for 10 min. Solubilized proteins (10 mg per condition) were incubated at least 2 hours with or without the TAT- conjugated peptide of interest before incubation overnight at 4 °C with immobilized C terminal peptide. Samples were washed twice with 5 ml of extraction buffer containing 500 mM Tris-EDTA, 250 mM NaCl (pH 8) and then twice with high sodium extracting buffer containing 500 mM Tris-EDTA, 500 mM NaCl (pH 8) before elution in SDS sample buffer. Samples were analyzed by Western blotting.

### Western Blot

Protein concentration in each lysate was determined by the bicinchoninic acid method. Proteins were resolved on 10% acrylamide gels and transferred electrophoretically onto nitrocellulose membranes (Amersham). Membranes were incubated in blocking buffer containing: PBS 0.1% Tween-20, and 5% skimmed dried milk for 1 hour at room temperature and overnight with primary antibodies in blocking buffer: chicken anti-PICK1 (1:500, NBP1-42829, Novus Biologicals), rat anti-HA (1:500, 11867423001, Roche) or monoclonal mouse anti-GFP antibody (1:1000, Clontech). Membranes were then washed and incubated with horseradish peroxidase-conjugated anti-chicken, anti-mouse or anti-rat secondary antibodies (1:4000 in blocking buffer, GE Healthcare) for 1 hour at room temperature. Immunoreactivity was detected with an enhanced chemiluminescence method (ECL detection reagent, GE Healthcare). Immunoreactive bands were quantified by densitometry using the Image J software (NIH).

### Endocytosis Experiments

Receptor internalization was evaluated using a fluorescence-based antibody-uptake assay described elsewhere [49]. Briefly, transfected neurons (12 DIV) were incubated with anti-HA antibody for 1 hour at 4°C to label surface-expressed mGlu3 receptors and then returned to conditioned media for 15 min at 37°C. Cells were fixed and incubated with biotin-conjugated first antibody and CF350-streptavidin (blue) to label the surface population of receptors. The cells were permeabilized and then incubated with Cy3-conjugated (red) secondary antibody to label the internalized population of receptors. The cells were mounted with Mowiol and imaged on a epifluorescent microscope equipped for optical sectioning (Apotome, Zeiss). Internalization was assessed in at least three independent experiments for each condition.

### Organotypic hippocampal slices

Organotypic hippocampal slices were prepared from 6-day-old wild-type (WT) C57Bl6/J mice as previously described [50] following a protocol approved by the veterinary Department of Animal Care of Montpellier. Three to five mice were used for each slice culture preparations and each experimental condition was tested over the course of at least three preparations. Slices were placed on a 30 mm porous membrane (Millipore, Billerica MA, USA) and kept in 100 mm diameter Petri dishes filled with 5 ml of culture medium containing 25% heat-inactivated horse serum, 25 % HBSS, 50% Opti-MEM, penicillin 25units/ml, streptomycin 25 μg/ml (Life technologies). Cultures were maintained in a humidified incubator at 37°C and 5% CO_2_ until DIV3 (day in vitro) and then were kept at 33 °C and 5% CO_2_ until the electrophysiological experiments.

### Electrophysiology

After 3 weeks in vitro, slices cultures were transferred to a recording chamber on an upright microscope (Olympus, France). Slices were superfused continuously at 1 ml/min with a solution containing (in mM) 125 NaCl, 2.7 KCl, 11.6 NaHCO_3_, 0.4 NaH_2_PO4, 1 MgCl_2_, 2 CaCl_2_, 5.6 D-glucose, and 0.001% phenol red (pH 7.4, osmolarity 305 mOsm) at 33°C. Whole cell recordings were obtained from cells held at - 70 mV using a Multiclamp 700B amplifier (Axon Instruments, Union City, CA, USA). Recording electrodes made of borosilicate glass had a resistance of 4-6 MΩ (Warner Instruments, USA) and were filled with (in mM) 125 K-gluconate, 5 KCl, 10 Hepes, 1 EGTA, 5 Na-phosphocreatine, 0.07 CaCl_2_, 2 Mg-ATP, 0.4 Na-GTP (pH 7.2, osmolarity 310 mOsm). Membrane potentials were corrected for junction potentials. For all recording conditions, only cells with access resistance < 18 MΩ and a change of resistance < 25% over the course of the experiment were analyzed. Data were filtered with a Hum Bug (Quest Scientific, Canada), digitized at 2 kHz (Digidata 1444A, Molecular Devices, Sunnyvale, CA, USA), and acquired using Clampex 10 software (Molecular Devices).

Activation of mGlu3 receptor by LCCG-1 (10 μM) induces an inward current in CA3 pyramidal cells (PCs) in the presence of TTX (1 μM), D-AP5 (40 μM) and picrotoxin (100 μM) [22].

### EEG/EMG recording and peptides administration

Wild-type C57BL6/J mice (10-12 weeks-old) were implanted for electroencephalographic (EEG) and electromyographic (EMG) recording and intra-ventricular peptide administration. All the procedures were conducted in accordance with the European Communities Council Directive (2010/63/EU). Mice were housed in groups of 5 per cage until surgery, and maintained in a 12 h light/dark cycle (lights on 7:30 am to 7:30 pm), in stable conditions of temperature (22± 2°C) and humidity (60%), with food and water provided *ad libitum*.

For surgery, animals were anesthetised with a mix of ketamine (100 mg/kg, Imalgene 500) and xylazine (10 mg/kg, Rompun 2%) plus a local subcutaneous injection of lidocaine (Xylocaine, AstraZeneca, France; 4 mg/kg in 50 μl of sterile 0.9% NaCl solution). They were then placed in a stereotaxic frame using the David Kopf mouse adaptor. A bipolar tungsten teflon-isolated torsade electrode was placed in the CA3 area of the left dorsal hippocampus (AP = −1.85, ML = +2.3, DV = −1.4 mm from bregma, https://scalablebrainatlas.incf.org). A 4 mm, 26 G canula guide (Phymep, France) was set into the left ventricle (AP = +0.1, ML = +0.75, DV = −2 mm from bregma). Skull cortical electrodes were placed on the frontoparietal bone and a reference electrode on the occipital bone. Two stainless steel EMG wires were placed in the neck muscles. All electrodes and the canula were fixed on the skull with dental acrylic cement. After surgery, mice were individually housed in order to minimize the chances of reciprocal injury caused by snatching of the implants. Freely moving animals were put into individual Plexiglass boxes, and their microconnectors were plugged to an EEG preamplifier circuit close to the head and to the EEG amplifier (Pinnacle Technology Inc., Lawrence, KS, USA). Mice were allowed 1 h 30 to habituate to the new environment and to the injection equipment (injection canula and tubing). TAT peptides were delivered via a pump operating a micro-syringe (injection volume: 5 μl, 500 μM in 0.9% NaCl, 750 nl / min). EEG data were obtained 30 min before and between 10 and 35 min post-injection. The electrical activity was recorded and filtered at 40 Hz, sampled at 200 Hz and recorded by a computer equipped with Sirenia® software (Pinnacle Technology Inc.). The electromyogram signal was filtered and sampled at 100 Hz. EEG recordings were performed together with video monitoring of the animal behavior. Sleep scoring and periodograms of the EEG recordings were obtained using Neuroscore (DSI, St. Paul, MN). Power spectral density was calculated using Fast-Fourrier Transform with 2 sec temporal and 0.5 Hz frequency resolutions.

### Statistical Data Analysis

Values are represented as mean ± SEM. Normality of data distribution was determined via the Shapiro-Wilk test. Except where noted, normally distributed data were analyzed via t-tests. For data sets with non-normal distributions, non-parametric tests were used (Mann-Whitney U test for independent samples or Wilcoxon signed-rank test for paired samples). Significance was defined as *P < 0.05, **P < 0.01,***P < 0.001.

Oscillation analyses were performed after 9 min of application of LCCG-1 or MCh. Calculation of oscillatory activity was performed from the time of the second peak in the Clampfit autocorrelation function. A segment was considered rhythmic when the second peak of the autocorrelation function was at least 0.3 and several regularly spaced peaks appeared.

## Acknowledgments

We specially thank Frederic De Bock for slice culturing, the animal facility (iExplore platform of RAM Montpellier, France) and Yan Chastagnier for technical assistance; Laurent Fagni, Philippe Marin and Emmanuel Valjent for valuable discussions; Corrado Corti, Corrado Corsi and GlaxoSmithKline for permission to use mGlu3 ^−/−^ mice.

